# Emotional and Linguistic Features Predict Prefrontal Functional Connectivity during Ongoing Dialogues: An fNIRS Investigation

**DOI:** 10.1101/2025.08.04.668428

**Authors:** Alessandro Carollo, Andrea Bizzego, Mengyu Lim, Massimo Stella, Gaia Doderovic, Gianluca Esposito

## Abstract

Identifying the neural bases of language has been a central focus in neuroscience since the pioneering case studies by Broca and Wernicke. Contemporary research has moved beyond classical modular models to conceptualize language as supported by a distributed network of anatomically and functionally interconnected brain regions. Yet, few studies have explored how these networks operate during spontaneous, real-life speech, limiting our understanding of language in natural contexts. In this study, we collected data from 84 individuals engaged in live conversations. Participants’ prefrontal brain activity was recorded using functional near-infrared spectroscopy hyperscanning, and functional connectivity was quantified via wavelet transform coherence. Dialogues were manually transcribed, and computational methods were applied to extract emotional and semantic/syntactic features from the speech data. Using linear mixed-effects models, we found that emotional content significantly predicted prefrontal functional connectivity 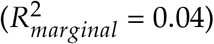 Among all emotional predictors, expressed anger was the most robust: higher anger levels were associated with reduced connectivity between the left middle frontal gyrus and the right inferior frontal gyrus 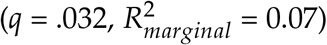. While other semantic and syntactic features did not predict overall connectivity, degree assortativity—an index of linguistic structure—was negatively associated with connectivity between the superior frontal gyri and the left inferior frontal gyrus 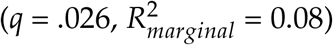. These findings highlight how both affective and structural properties of speech modulate prefrontal connectivity during real-life interaction. More broadly, this work demonstrates the potential of integrating computational linguistics with social neuroscience to uncover the neural mechanisms of real-life social interactions.

## 1. Introduction

Language is the primary medium through which human beings connect with one another. It enables people to convey emotions, share thoughts, transmit knowledge, build social relationships, and organize complex cultural and institutional structures. In everyday life, language affects a variety of activities, from interpersonal interactions to more complex cognitive processes (Carollo et al., 2025b; Tomasello, 2010). Given its pervasive role in human behavior and cognition, understanding the neural basis of language has become a central goal in cognitive neuroscience.

Since the earliest neurological case studies (Broca, 1861; Geschwind, 1970; Lichtheim et al., 1885; Wernicke, 1969), the neuroscience of language has progressively evolved, providing more fine-grained descriptions of how language is encoded in and supported by the brain (Tremblay and Dick, 2016). The classical model of language originates from the clinical observations of Paul Broca and Carl Wernicke, who were among the first to link specific brain regions to distinct linguistic and behavioral outcomes. Particularly, Broca associated the left posterior inferior frontal gyrus with language articulation and syntactic processing. In contrast, Wernicke identified common lesions in the superior temporal gyrus in patients who had difficulty comprehending spoken language, despite relatively preserved articulation abilities. Subsequently, the work by Lichtheim et al. (1885) expanded this model by proposing neuroanatomical pathways responsible for mediating communication between these functionally specialized regions. However, the classical model’s emphasis on discrete, modular areas with narrowly defined roles has since been challenged, as growing evidence supports a more distributed and interactive network underlying language processing (Friederici et al., 2017).

Building on this paradigm shift, more recent studies have moved beyond the modular approach of the classical model to describe language as emerging from a network of anatomically and functionally interconnected brain regions (Poeppel et al., 2012; Price, 2012). From this perspective, the language network involves multiple interconnected pathways, including fronto-temporal, parieto-temporal, occipito-temporal, and fronto-frontal connections, as well as cortico-subcortical loops (Tremblay and Dick, 2016). For instance, the dual stream model of language has received wide attention in the scientific literature (Hickok and Poeppel, 2007; Rauschecker and Scott, 2009). This model identifies two main streams of language processing in the brain: the dorsal and ventral pathways. The dorsal pathway includes fronto-temporo-parietal regions involved in mapping auditory speech sounds to articulatory representations (Dick et al., 2014). In parallel, the ventral pathway is involved in semantic comprehension and in processing less complex syntactic structures (Dick et al., 2014). However, as discussed by Grappe et al. (2019), much of the evidence supporting network-based models of language comes from comparative research in macaque monkeys and, in humans, from structural connectivity analyses using diffusion tensor imaging and functional connectivity estimates derived from resting-state functional magnetic resonance imaging (fMRI). Relatively few studies have directly investigated how these networks function during active language production (e.g., Ewald et al., 2012; Grappe et al., 2019), leaving a significant gap in our understanding of real-time neural dynamics underlying ongoing linguistic processing.

In the present work, we aim to bridge this gap by employing functional near-infrared spectroscopy (fNIRS) to monitor brain activity and assess prefrontal functional connectivity during active, real-life social interactions (Carollo et al., 2025b; Lim et al., 2024a,b). Compared to traditional neuroimaging techniques such as electroencephalography (EEG) and fMRI, fNIRS is less susceptible to motion artifacts, making it well-suited for studying dynamic and ecologically valid social interactions without compromising signal quality (Bizzego et al., 2025, 2024). Building on these properties, the current study seeks to explore how real-life language use modulates prefrontal connectivity, with particular attention to the emotional and linguistic features of spoken interaction. To achieve this, we leverage recent advances in affective computing and cognitive network modeling (Carollo et al., 2025b). Specifically, we employ EmoAtlas to extract quantitative measures of the emotional content embedded in speech, and *Textual Forma Mentis Networks* (TFMNs) to map the conceptual associations within participants’ discourse (Semeraro et al., 2025; Stella et al., 2019).

## 2. Methods

### 2.1. Study design

The dataset analyzed in this study originates from a cross-cultural project investigating the neural mechanisms underlying role-play, a therapeutic technique commonly used to address psychopathological symptoms (Lim et al., 2024a,Lim et al., 2024b,Lim et al., 2024c). Brain activity was measured using a hyperscanning protocol based on fNIRS, while dyads of participants engaged in three distinct five-minute interaction tasks: natural conversation, role-play, and role reversal. Although the study was conducted in both Italy and Singapore, only the Italian cohort was included in the current analysis. This decision was made to reduce linguistic variability, as participants in Singapore often used multiple languages and dialectal forms such as Singlish—a creole language not compatible with the computational tools employed for linguistic analysis (Carollo et al., 2025b).

All interactions were video-recorded and subsequently transcribed. To extract features from the dialogues, we applied two automated tools: EmoAtlas (Semeraro et al., 2025), which quantifies the emotional content of speech, and TFMNs (Stella et al., 2019), which capture the underlying syntactic and semantic relationships. The resulting emotional *z*-scores and TFMNs network properties were then used to model functional connectivity patterns across specific subregions of the prefrontal cortex. Figure 1 provides a graphical summary of the study design.

**Figure 1:**
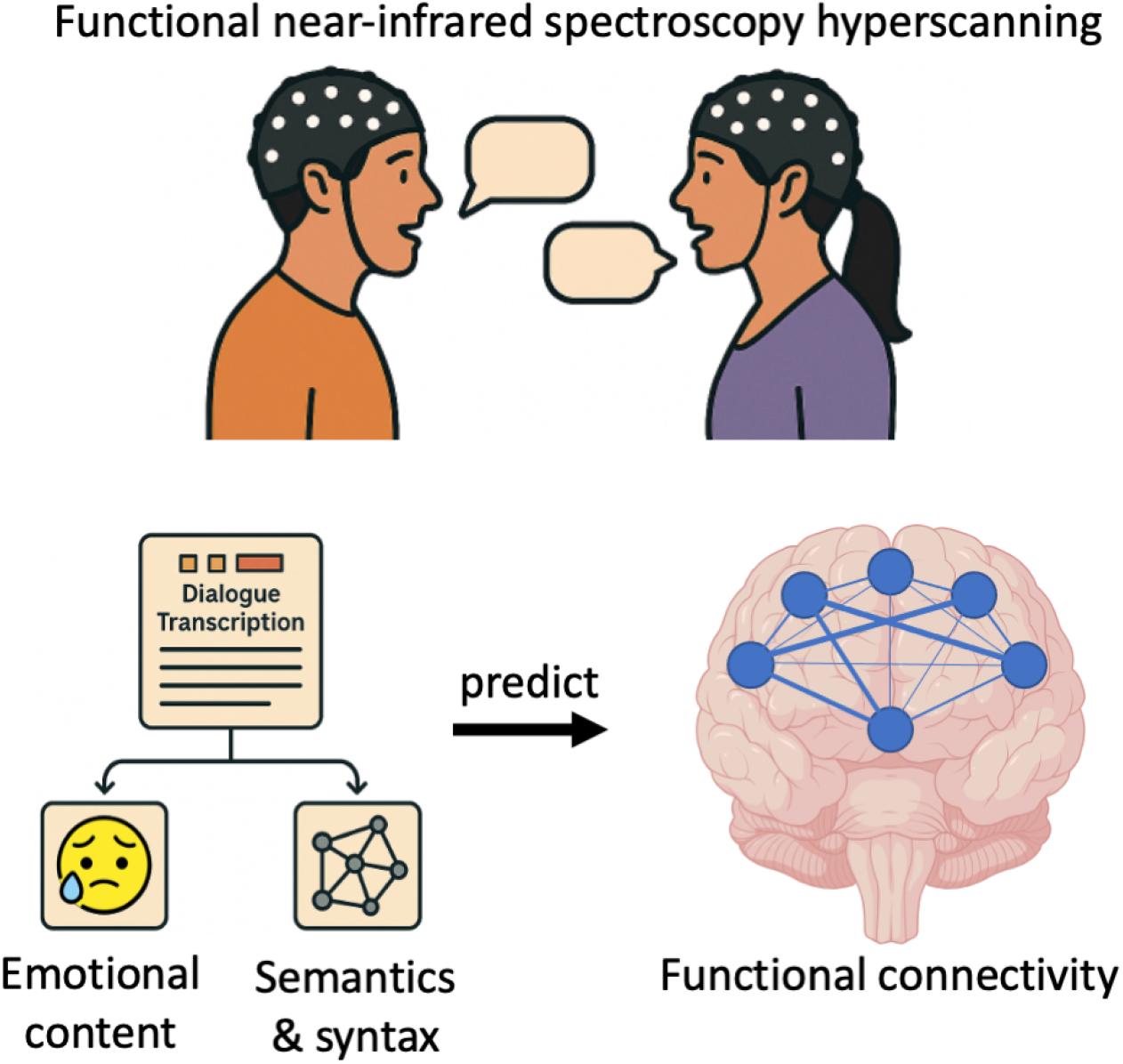
Graphical representation of the study design. Participants took part in a functional near-infrared spectroscopy hyperscanning session while engaging in verbal conversations. The recorded dialogues were manually transcribed and analyzed using computational methods to extract both emotional content and semantic/syntactic features. These emotional and linguistic properties were then used as predictors of prefrontal functional connectivity, assessed through wavelet transform coherence.

Ethical approval for data collection was obtained from both the University of Trento (2022-059) and Nanyang Technological University (NTU-IRB-2021-03-013). The entire study was conducted in accordance with the Declaration of Helsinki, and informed consent was secured from all participants prior to their involvement.

### 2.2. Participants

A total of 84 individuals (forming 42 dyads; age range: 18–35 years) took part in the study. Participants were recruited through convenience and snowball sampling via social media platforms. Each dyad was composed of peers who already shared an existing friendship. None of the participants reported a history of neurological or medical conditions that could interfere with cerebral oxygenation, such as disorders affecting hemoglobin function. Furthermore, all participants confirmed they were not taking medications known to influence cerebral blood flow or systemic blood pressure at the time of participation.

### 2.3. Experimental protocol

The experimental protocol was structured into four distinct phases, during which neural activity was recorded using fNIRS (Lim et al., 2024a,Lim et al., 2024b,Lim et al., 2024c). The session started with a two-minute resting-state baseline, during which participants sat quietly facing each other while minimizing limb movement to establish a reference of neural activity at rest. Following this, participants engaged in three five-minute interactive conditions: *(1)* natural conversation, *(2)* role-play, and *(3)* role reversal.

During the natural conversation phase, participants were encouraged to interact naturally, as they would in everyday situations. In the role-play condition, each participant assumed the identity of a mutual friend from their peer group, simulating an imaginary conversation. In the role reversal condition, the two participants exchanged roles, with each one impersonating the other. All interactive conditions were guided by a shared narrative prompt: participants were asked to imagine unexpectedly meeting at a shopping mall while shopping for each other’s gifts. To control for order effects, the sequence of the three interaction phases was counterbalanced across dyads.

In line with prior work (Levinson, 1983; Lim et al., 2024c), the initial and final minutes of each five-minute interaction were excluded from analysis to minimize variability introduced by conversation openings and closings. The resting-state data were also excluded from the present study, as the primary focus was on language production phases to explore how emotional and syntactic/semantic properties of conversations relate to functional connectivity within the frontal cortex.

### 2.4. Acquisition of neural data

During the experiment, participants’ brain activity was recorded with an fNIRS hyperscanning approach while dyads engaged in social interactions. To record neural activity from the prefrontal cortex, participants wore fNIRS caps equipped with 8 light sources and 7 detectors, configured in accordance with a standard prefrontal montage (Azhari et al., 2019; Carollo et al., 2025b). Each source emitted near-infrared light at two wave-lengths—760 nm and 850 nm—while the detectors captured the reflected signals, resulting in 20 functional channels per participant. These channels were spatially organized to monitor key regions within the prefrontal cortex, which were grouped into six regions of interest: anterior prefrontal cortex, superior frontal gyri, left and right middle frontal gyrus, and left and right inferior frontal gyrus. Because of their central location and the spatial constraints of fNIRS technology, the anterior prefrontal cortex and superior frontal gyri were not further divided into hemispheric subregions.

The optode arrangement followed the international 10–20 system commonly used in EEG studies (Homan et al., 1987), ensuring standardized placement across participants. To maintain optimal signal quality, the distance between each light source and detector was kept below 3 cm, using optode stabilizers to preserve a high signal-to-noise ratio (Pinti et al., 2020). Data acquisition was carried out using a NIRSport2 system (NIRx Medical Technologies LLC), with a sampling frequency of 10.17 Hz.

### 2.5. Processing of neural data

Neural data preprocessing was carried out using the *pyphysio* library (Bizzego et al., 2019). To evaluate signal quality, we employed a deep learning-based approach, specifically a convolutional neural network trained to classify fNIRS signal segments according to their quality (Bizzego et al., 2022b, 2021). Motion-related artifacts were corrected using a combination of spline interpolation (Scholkmann et al., 2010) and wavelet filtering techniques (Molavi and Dumont, 2012).

Following artifact correction, fNIRS signals were transformed into concentrations of oxygenated (HbO) and deoxygenated (HbR) hemoglobin using the modified Beer–Lambert law (Delpy and Cope, 1997). Consistent with prior literature (e.g., Carollo et al., 2025a,Carollo et al., 2025b), our main analyses focused on the HbO component.

To extract regional neural activity, signal values from individual channels were grouped according to predefined anatomical regions of interest. For each region of interest, we computed the average signal across normalized channels, provided that at least two good-quality channels were available within the region (Bizzego et al., 2022a).

### 2.6. Functional connectivity

To assess functional connectivity within participants’ prefrontal cortex, we computed wavelet transform coherence (WTC) between time series from different regions of interest (Chang and Glover, 2010; Grinsted et al., 2004). This method allows for the examination of coherence between neural signals as a function of both time and frequency, capturing both in-phase and phase-lagged relationships across signals and providing a comprehensive view of functional interactions between cortical regions.

WTC was applied to each region pair for each participant and experimental condition (i.e., natural conversation, role-play, and role reversal), resulting in a total of 15 WTC values per participant. Coherence was computed across a frequency range from 0.01 to 0.20 Hz, with a step size of 0.01 Hz (Carollo et al., 2025a,b). The final connectivity metric for each combination of regions of interest and experimental condition was obtained by averaging coherence values across all frequency bands, as no specific frequency hypotheses were posited *a priori* (Carollo et al., 2025a,Carollo et al., 2025b).

### 2.7. Emotional content of dialogues

To examine the emotional aspects present in the transcribed conversations, we utilized EmoAtlas (Semeraro et al., 2025), a tool designed to construct TFMNs and perform emotional profiling based on Plutchik’s emotion theory (Plutchik, 1980). EmoAtlas quantifies eight fundamental emotions within text: anger, anticipation, disgust, fear, joy, sadness, surprise, and trust. For each emotion, it generates a *z*-score reflecting the relative strength of that emotion in the dialogue compared to randomly assembled word sets drawn from its psychological lexicon.

Specifically, given a text with *m* emotional words, random word sets of the same size are repeatedly sampled uniformly from EmoAtlas’s emotional vocabulary. Because the sampling is uniform, the random sets naturally incorporate the frequency biases of the lexicon (e.g., words associated with trust appear more often than those linked to disgust). This random sampling process is repeated 1000 times to create a distribution of counts 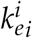 where 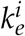 represents the number of words related to emotion *e* in the *i*-th random sample. The emotional *z*-score for each emotion is then calculated by comparing the observed number of words *m*_*e*_ in the actual text to the mean and standard deviation of the random samples:

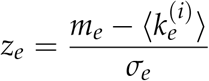

where *σ*_*e*_ is the standard error of the mean across the random samples. EmoAtlas was selected based on its demonstrated accuracy, which matches or exceeds that of leading natural language processing methods (Semeraro et al., 2025).

### 2.8. Syntactic/semantic structure of dialogues

To examine the syntactic and semantic organization of the dialogues, we utilized TFMNs (Stella et al., 2019), constructed via EmoAtlas (Semeraro et al., 2025). TFMNs represent the associative structure of language by modeling words or concepts as nodes connected through syntactic (grammatical relations) or semantic (synonym) links. The method involves segmenting the text into sentences and, within each sentence, connecting words that are within a syntactic distance of *K* ≤ 4 on the sentence’s parsing tree. This approach allows TFMNs to capture syntactic dependencies even between words that are not immediately adjacent, a capability that surpasses traditional word co-occurrence networks (Quispe et al., 2021). The syntactic parsing trees are generated using a pre-trained artificial intelligence model (from *spaCy*; see Semeraro et al. 2025), analyzing each sentence separately. The distance threshold *K* was set to 4 based on previous research indicating that English syntactic distances generally average around 3, reflecting linguistic efficiency principles (i Cancho et al., 2004). After identifying syntactic links among non-stopwords within this threshold, the network is enriched with semantic relationships and psychological-emotional annotations, resulting in a multiplex network where edges represent either syntactic or semantic connections and nodes are tagged with emotional valence (positive, negative, neutral) and emotion categories (Semeraro et al., 2025, 2022; Stella et al., 2019).

After constructing the semantic network for each participant across all conditions, we calculated several network metrics (Siew et al., 2019; Stella, 2022): total number of nodes |*V*|, total number of edges |*E*|, average local clustering coefficient, number of connected components |*C*|, and degree assortativity. The total number of nodes |*V*| represents the unique words or concepts included in the network, while the total number of edges |*E*| reflects the syntactic or semantic connections between them. For example, in a semantic network derived from a text, |*V*| corresponds to the count of distinct words or concepts, and |*E*| denotes how these terms are linked based on their syntactic or semantic relationships. The average local clustering coefficient measures the probability that two neighbors of a node are also connected, indicating the tendency for nodes to form tightly-knit groups. In the context of semantic networks, a higher clustering coefficient suggests that concepts are densely organized into localized clusters. The number of connected components |*C*| identifies how many disconnected subgroups exist within the network, where each subgroup is internally connected but isolated from others. For instance, if the network breaks into four distinct groups of terms with no inter-group links, then |*C*| equals 4. Lastly, degree assortativity quantifies the tendency of nodes to connect with others having a similar number of connections (i.e., degree); higher values indicate that highly connected nodes tend to link with other highly connected nodes.

### 2.9. Data analysis

For the statistical analyses in the current study, we used linear mixedeffects models (Bates et al., 2005), which were selected because they account for both fixed and random effects, allowing us to appropriately model the nested structure of our data. In particular, we divided the statistical plan as follows.

First, we conducted an investigation of the relationship between emotional content of speech and prefrontal functional connectivity. We first conducted a linear mixed-effects model with functional connectivity scores as the dependent variable and emotional *z*-scores as fixed effects. The model included region of interest combination ID, experimental condition, dyad ID, and participant ID as random effects. To further investigate the relationship between emotional content and functional connectivity at the region of interest level, we ran separate linear mixed-effects models for each region of interest pair. In each of these models, functional connectivity was the dependent variable, emotional *z*-scores were included as fixed effects, and experimental condition, dyad ID, and participant ID were modeled as random effects. To account for multiple comparisons, we applied the Benjamini-Hochberg correction to adjust the alpha level (Benjamini and Hochberg, 2000).

Second, we conducted an investigation of the relationship between semantic and syntactic properties of speech and prefrontal functional connectivity. We applied the same analytical strategy described above, this time including the TFMNs metrics as fixed effects.

## 3. Results

### 3.1. Emotional content of speech and functional connectivity

In the first part of our analysis, we examined the relationship between the emotional content of speech and functional connectivity in the prefrontal cortex.

We first ran a linear mixed-effects model using data from the entire prefrontal cortex, with functional connectivity scores as the dependent variable and the emotional content of speech as fixed effects. The model revealed that higher levels of anger (*β* = –0.030, *p <*.001) and anticipation (*β* = –0.012, *p* =.002) were significantly associated with lower prefrontal functional connectivity. In contrast, higher levels of surprise (*β* = 0.016, *p <*.001), disgust (*β* = 0.009, *p* =.034), and sadness (*β* = 0.011, *p* =.005) significantly predicted increased functional connectivity. Trust, joy, and fear did not emerge as significant predictors of functional connectivity (*p*s *>*.050). The model explained a small but meaningful portion of the variance, with a marginal *R*^2^ of 0.03. The full results of this model are reported in Table 1.

**Table 1:**
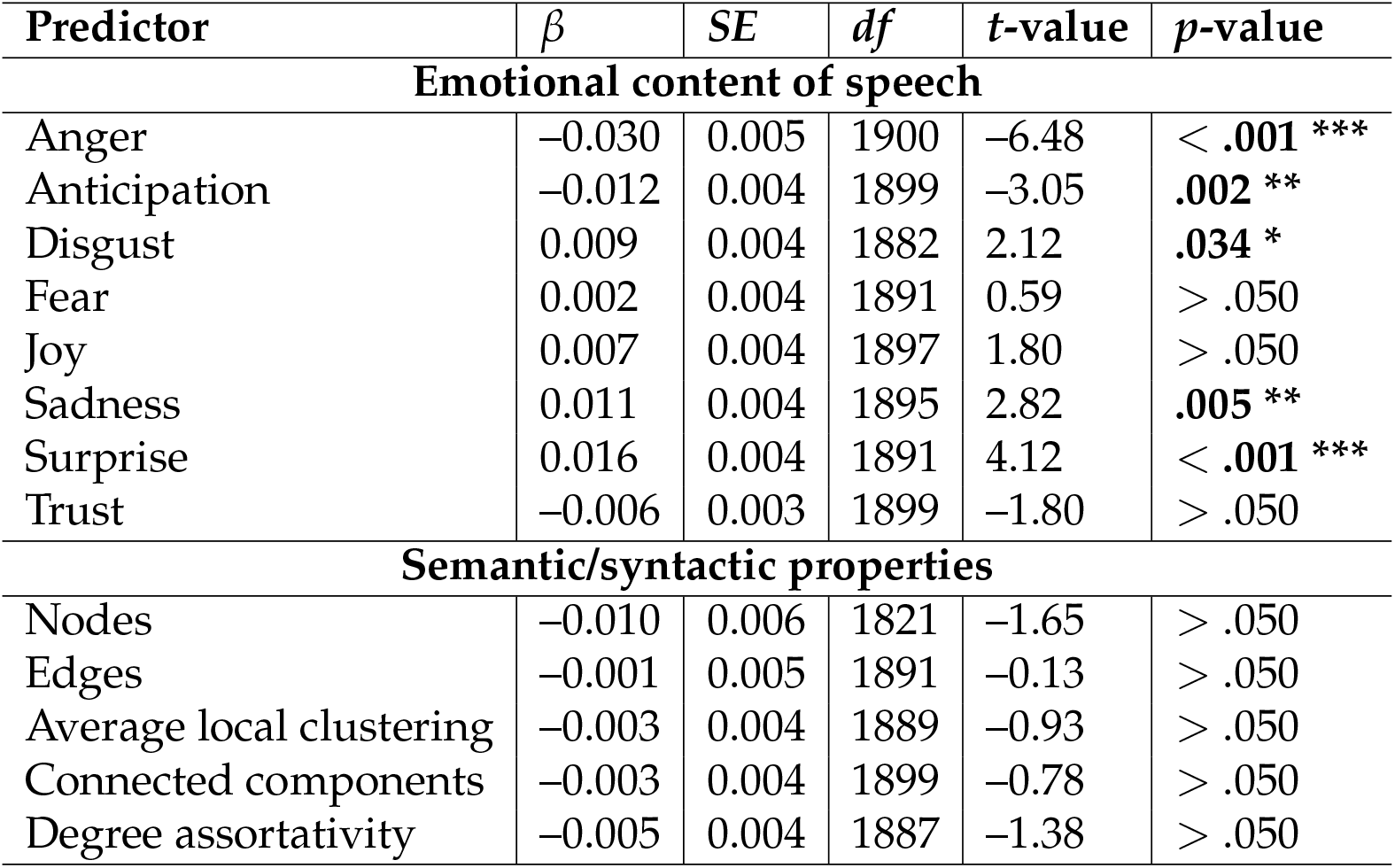
Results of the linear mixed-effects models predicting functional connectivity patterns across the whole prefrontal cortex. For each predictor, we report the estimate (*β*), the standard error (*SE*), the degrees of freedom (*df*), the *t*-value, and the *p*-value. (^*^ *p <*.05; ^**^ *p* <.01; ^***^ *p* <.001).

We then ran separate linear mixed-effects models for each pairwise combination of regions of interest, analyzing functional connectivity patterns individually. After adjusting the *p*-values using the Benjamini-Hochberg correction, we found a statistically significant negative association between anger and functional connectivity between the anterior prefrontal cortex and the right middle frontal gyrus (*β* = –0.045, *p* =.006, *q* =.049), as well as between the left middle frontal gyrus and the right inferior frontal gyrus (*β* = –0.035, *p* =.004, *q* =.032). These two models had a marginal *R*^2^ of 0.08 and 0.07, respectively. None of the other emotional *z*-scores significantly predicted functional connectivity across regions of interest (*q*s >.050). The details regarding statistically significant predictors of these models are reported in Table 2.

**Table 2:** Statistically significant predictors of functional connectivity patterns across combinations of regions of interest. For each predictor, we report the estimate (*β*), the standard error (*SE*), the degrees of freedom (*df*), the *t*-value, the *p*-value, and the FDR-corrected *q*-value. Abbreviations: aPFC = anterior prefrontal cortex, IFG = inferior frontal gyrus, MFG = middle frontal gyrus, SFG = superior frontal gyri. (^*^ *q* <.05).

| Regions of interest | Predictor | $\beta$ | SE | df | t-value | p-value | q-value |
| --- | --- | --- | --- | --- | --- | --- | --- |
| **Emotional content** |  |  |  |  |  |  |  |
| aPFC – right MFG | Anger | -0.045 | 0.016 | 92.60 | -2.80 | .006 | **.049 \*** |
| left MFG – right IFG | Anger | -0.035 | 0.012 | 101.29 | -2.97 | .004 | **.032 \*** |
| **Semantic/syntactic properties** |  |  |  |  |  |  |  |
| SFG – left IFG | Degree assortativity | -0.041 | 0.014 | 94.57 | -2.90 | .004 | **.026 \*** |

### 3.2. Semantic/syntactic structure of speech and functional connectivity

In the second part of our analysis, we examined whether the semantic/syntactic structure of speech was associated with changes in prefrontal functional connectivity.

We began by running a linear mixed-effects model on the full prefrontal cortex dataset, using functional connectivity scores as the dependent variable and the TFMNs metrics as fixed effects. The results indicated that none of the TFMNs metrics were significantly associated with functional connectivity across the prefrontal cortex (*p*s *>*.050). A detailed summary of the model results is provided in Table 1.

We then conducted individual linear mixed-effects models for each pairwise combination of regions of interest to examine localized patterns of functional connectivity. Following Benjamini-Hochberg correction for multiple comparisons, we identified a statistically significant negative association between degree assortativity index and the functional connectivity between the superior frontal gyri and the left inferior frontal gyrus (*β* = –0.041, *p* =.004, *q* =.026). In other words, speech characterized by more common words being connected to less frequent ones was associated with higher connectivity between the superior frontal gyri and the left inferior frontal gyrus. This model had a marginal *R*^2^ of 0.08. None of the other TFMNs metrics significantly predicted functional connectivity across regions of interest (*q*s >.050). The details regarding statistically significant predictors of these models are reported in Table 2.

## 4. Discussion

In the present study, we investigated how the emotional content and the semantic/syntactic features of naturalistic speech relate to patterns of prefrontal functional connectivity during real-time social interactions. To address this question, we employed an integrative computational framework that combines quantitative linguistic analysis with advanced neuroimaging techniques, enabling a fine-grained examination of the neural correlates of spoken language in ecologically valid settings.

Our findings suggest that the emotional content of speech is a stronger predictor of overall prefrontal functional connectivity than its semantic or syntactic properties. The prefrontal cortex regions covered by our setup encompass areas that functionally correspond to both the dorsolateral and ventrolateral prefrontal cortices. Both these regions, as well as their functional connectivity, have a prominent role in emotion regulation, particularly in the implementation of cognitive reappraisal strategies (Li et al., 2022; Sanchez-Lopez et al., 2018; Tang et al., 2025). In contrast, semantic and syntactic processing could rely on more localized brain regions within the prefrontal cortex. As a result, including all regions of interest in our connectivity analyses may have introduced variability unrelated to these specific linguistic processes, potentially reducing the strength of their predictive power.

With regard to the emotional content of speech, expressed anger emerged as a robust and consistent negative predictor of functional connectivity, particularly between the left middle frontal gyrus and the right inferior frontal gyrus. Specifically, our analysis showed that participants who used anger-related words more frequently exhibited reduced connectivity between these two regions. Both the right inferior frontal gyrus and the left middle frontal gyrus show atypical patterns of activation and deactivation in individuals with psychiatric conditions marked by emotional dysregulation (e.g., schizophrenia, depression, borderline personality disorder; Engels et al., 2010; Pilon et al., 2025; Xiao et al., 2024). In the meta-analysis by Sorella et al. (2021), the right inferior frontal gyrus was identified as a region consistently activated during both the perception and the experience of anger. The middle frontal gyrus, or dorsolateral prefrontal cortex, is known to play a key role in the top-down regulation of emotional attention, with greater excitability in the left hemisphere being associated with faster disengagement from emotional stimuli (Alizadehgoradel et al., 2024; De Raedt et al., 2010; Sanchez-Lopez et al., 2018). Thus, the expression of anger in speech may be accompanied, and possibly sustained, by a functional disconnection between a region implicated in anger perception and experience (the right inferior frontal gyrus) and a region involved in attentional disengagement toward emotional stimuli (the left middle frontal gyrus). On the one hand, the simultaneous activation of these two regions could possibly underlie the inhibition of anger expression and attentional disengagement from emotional content in speech. On the other hand, functional disconnection could allow for the attentional engagement and consequent expression of anger-related words.

With regards to the semantic/syntactic properties of language, linguistic complexity, captured by lower values of the degree assortativity index, was a significant predictor of functional connectivity between the superior frontal gyri and the left inferior frontal gyrus. Specifically, we observed that greater semantic/syntactic complexity and unpredictability (i.e., lower degree assortativity values) was associated with stronger connectivity between these two regions. Both the left inferior frontal gyrus and the superior frontal gyri are key regions involved in the processing of linguistic information. The left inferior frontal gyrus, often linked to Broca’s area, plays a crucial role in the selection of semantic knowledge and is actively engaged in syntactic computations during both speech production and comprehension (Fedorenko et al., 2024; Giglio et al., 2022; Thompson-Schill et al., 1997; Tyler et al., 2011). Additionally, the left inferior frontal gyrus is sensitive to linguistic complexity and structural hierarchies (Friederici et al., 2006; Yang et al., 2017). The superior frontal gyri, on the other hand, are implicated in a variety of cognitive and motor control tasks (Li et al., 2013). With regard to semantics, the study by Sharp et al. (2010) demonstrated that the left prefrontal cortex is more strongly activated during decisions involving semantic, as opposed to perceptual, complexity. Among the regions showing sensitivity to semantic complexity is the left superior frontal gyrus. Notably, there is extensive structural connectivity between the left inferior frontal gyrus and the superior frontal gyri, among which is the frontal aslant tract (Bohsali et al., 2025; Catani et al., 2013; Dick et al., 2019). Overall, this connectivity is thought to serve as the neural substrate for the coordination of syntax and grammatical morphology as well as for the planning, timing, and coordination of sequential motor movements (Bohsali et al., 2025; Catani et al., 2013; Dick et al., 2019). These findings align with our results, which show that greater linguistic complexity is associated with increased functional connectivity between the left inferior frontal gyrus and the superior frontal gyri. This suggests that as linguistic production becomes more semantically and syntactically demanding, there is a heightened need for dynamic coordination between these regions.

## 5. Limitations

Some aspects should be acknowledged as potential limitations of the current work and could potentially inspire future lines of research in the field.

Specifically, this study used fNIRS to investigate brain activity only in the prefrontal cortex. While this region is known to play a key role in both linguistic and emotional processing, exploring additional brain areas not covered by our cap setup could offer further insight into how language properties shape functional connectivity at the neural level. Moreover, in this experiment, fNIRS was chosen because of its greater tolerance to head movements compared to other neuroimaging techniques, which makes it particularly suitable for implementing tasks such as live conversations without compromising signal quality. Nonetheless, other methodologies, such as EEG, could provide a more fine-grained understanding of the temporal patterns of functional connectivity.

## 6. Conclusions

In the present work, we integrated an affective and cognitive computational framework for language analysis (Semeraro et al., 2025; Stella et al., 2019) with recent advances in social and cognitive neuroscience (Carollo et al., 2025b) to investigate how the emotional and linguistic properties of spontaneous speech relate to functional connectivity within the prefrontal cortex. Our results indicate that the emotional content of language is a robust predictor of prefrontal connectivity. In particular, higher levels of expressed anger are significantly associated with reduced connectivity between the left middle frontal gyrus and the right inferior frontal gyrus, suggesting a transient disconnection between these regions during emotionally charged moments. Conversely, lower levels of anger are linked to more integrated connectivity, reflecting a more cohesive prefrontal network under calmer affective states. In addition, we found that greater linguistic complexity and unpredictability are associated with stronger connectivity between the superior frontal gyri and the left inferior frontal gyrus. This pattern may reflect the engagement of these regions in the generation of more complex speech. Overall, we observe a functional dissociation in prefrontal connectivity, whereby central-left connectivity is more strongly associated with syntactic and semantic properties of speech, while central-right connectivity is more closely linked to its emotional content. These results underscore the role of affective and semantic/syntactic components in shaping the neural mechanisms that support spontaneous speech production and provide new insights into how functional neuroimaging can be used to explore the brain dynamics during ongoing social interaction.

## Author contributions

Conceptualization: AC, GE; Data curation: AC, ML; Methodology: AC, MS, AB; Formal Analysis: AC; Investigation: AC, ML; Writing–original draft preparation: AC, GD; Writing–review and editing: AC, MS, AB, ML, GD, GE; Supervision: GE. All authors have read and agreed to the published version of the manuscript.

